# Voice patches in the marmoset auditory cortex revealed by wide-field calcium imaging

**DOI:** 10.1101/2024.02.19.581089

**Authors:** Yang Zhang, Xindong Song, Yueqi Guo, Chenggang Chen, Michael S Osmanski, Xiaoqin Wang

**Affiliations:** Laboratory of Auditory Neurophysiology, Department of Biomedical Engineering, Johns Hopkins University School of Medicine, Baltimore, Maryland 21205, USA

## Abstract

Species-specific vocalizations are behaviorally critical sounds. Similar to faces, species-specific vocalizations are important for the survival and social interactions of both humans and vocal animals. Face patches have been found in the brains of both human and non-human primates. In humans, a voice patch system has been identified on the lateral superior temporal gurus (STG) that is selective to human voices over other sounds. In non-human primates, while vocalization-selective regions were found on the rostral portion of the temporal lobe outside of the auditory cortex in both macaques and marmosets using functional magnetic resonance imaging (fMRI), it is yet clear whether vocalization-selective regions are present in the auditory cortex. Using wide-field calcium imaging, a technique with both high temporal and high spatial resolution, we discovered two voice patches in the marmoset auditory cortex that preferentially respond to marmoset vocalizations over other sounds and carry call types and identity information. One patch is located on the posterior primary auditory cortex (A1), and the other one is located on the anterior non-core region of the auditory cortex. These voice patches are functionally connected and hierarchically organized as shown by latency and selectivity analyses. Our findings reveal the existence of voice patches in the auditory cortex of marmosets and support the notion that similar cortical architectures are adapted for recognizing communication signals for both vocalizations and faces in different primate species.

## INTRODUCTION

The human voice is the most critical sound of our daily life which carries a wealth of social information. It not only contains speech information, but also can be viewed as an “auditory face” that allows us to recognize individuals and their emotional states. The ability to analyze and categorize the information contained in human voice plays a key role in human social interactions. Because of the uniqueness of the human voice, scientists are wondering whether the human brain has specialized regions for processing the human voice. Functional imaging studies in humans have demonstrated the existence of voice-specific regions, which are located on the lateral superior temporal gurus (STG) and in the upper bank of the superior temporal sulcus (STS)^1–4^. These regions prefer human voices over other sounds including acoustic controls. Further studies suggest a voice patch system, which consists of a series of discrete and interconnected cortical regions on the STG, is specialized for processing human voices in the human brain^5–6^. These findings provide evidence indicating that there are specialized regions in the human brain dedicated to processing high biological relevant stimuli such as human voices.

In non-human primates, species-specific vocalizations are behaviorally critical sounds. Many non-human primates readily use species-specific vocalizations for communications. For example, marmosets can take turns back and forth to exchange their vocalizations with conspecifics (antiphonal calling) just as in human conversations^7–10^. In addition, non-human primates can recognize conspecifics by their vocalizations^11–12^. Analog to the face-selective patches that have been found in both humans and non-human primates^13–15^, the auditory system of non-human primates might have also evloved specialized regions dedicated to processing species-specific vocalizations. Functional imaging studies in macaques^16^ and marmosets^17^ suggest the existence of vocalization-specific regions in the brain, which are both located on the rostral portion of the temporal lobe outside the auditory cortex. These findings reveal that the voice/vocalization-specific regions are evolutionarily conserved in primates.

In contrast to the voice patch system found in humans specialized for processing human voice, functional magnetic resonance imaging (fMRI) of non-human primates^16–17^ provides evidence predominantly for one concentrated vocalization-specific region outside the auditory cortex. There has never been a description of a voice patch system consisting of a series of discrete and interconnected cortical regions analogous to the human voice patch system in non-human primates. In addition, using fMRI to investigate auditory functions is very challenging due to the substantial acoustic noises generated by fMRI sequences^18^. Although sparse sampling imaging technique^19^, which allows stimuli to be presented in quiet and the peak BOLD (blood oxygen level dependent) activity to be captured due to the delay in hemodynamic responses, has been widely used in auditory neuroscience research^16–17^, it is at a cost of reduced temporal resolution. Furthermore, compared with neurophysiology signals which directly reflect signals from neurons, fMRI BOLD signals have a relatively lower signal-to-noise ratio (SNR). Therefore, a technique with high temporal and spatial resolution as well as high SNR is needed to investigate whether there are voice patches in the auditory cortex of non-human primates.

Here, we used wide-field calcium imaging in marmosets to tackle this problem. The marmoset is an increasingly popular biomedical non-human primate model^20–23^. It is a highly vocal and social primate species with a sophisticated vocal repertoire for vocal communication^24–25^. It has become an ideal animal model to fill the translational gap between rodent models and humans^26–27^. By implanting a chronic cranial window covering the entire marmoset auditory cortex and injecting the virus with calcium indicator into the cortex through the artificial dura, we were able to record the calcium signals from the entire auditory cortex while presenting the species-specific vocalizations and other acoustic controls to the awake marmosets. We identified two voice patches in the marmoset auditory cortex in each hemisphere. The voice patches were hierarchically organized and functionally connected. Moreover, call types and identity information are carried by population responses from the voice patches.

## RESULTS

### Voice patches in marmoset auditory cortex

To record calcium signals in awake marmosets, we implanted a cranial window with artificial dura over the temporal lobe that covers the entire auditory cortex based on the tonotopy maps obtained by through-skull imaging of intrinsic signals as reported previously^28^. Fig. 1a illustrates the position of a typical cranial window in this study. Viruses with calcium indicators (GCaMP6) were injected into the cortex through the artificial dura in two marmosets (Fig. 1b, left hemisphere for M56E, and right hemisphere for M160E) using the injection techniques developed in our lab^29^. Fig. 1b shows the GCaMP6 expression levels in the two marmosets (brighter signals indicate higher expression levels). The borders of the cranial windows are marked by white dashed lines. To show the overall organization of the auditory cortex, we first obtained the tonotopic maps within the cranial window based on intrinsic signals using a phase-encoded Fourier approach^30–31^ as described in our recent publication^28^. Fig. 1c shows the tonotopic gradients of the auditory cortex in two marmosets (red: low-frequency, purple: high-frequency). In each animal, the white dotted line indicates the tentative border between core (tone responsive) and non-core regions of the auditory cortex. To locate cortical regions that may be selectively responsive to marmoset vocalizations, we recorded calcium signals within the cranial windows while seven categories of sounds were presented to the marmosets. The seven categories of sounds are marmoset vocalizations (MarV), macaque vocalizations (MacV), human speech (HumS), other animal vocalizations (AmV), natural sounds (NS), artificial sounds (AS), and scrambled marmoset vocalizations (SMarV). Marmosets have a rich vocal repertoire for vocal communications^24–25^. Several distinct classes of vocalizations have been described for this species^32^. We used the 4 most commonly produced call types (Phee, Trillphee, Trill, and Twitter) recorded from 6 marmosets in this study (6 samples for each call type from each marmoset). Extended Data Fig. 1 shows the exemplars of the four call types which have distinct acoustic features. We then compared the responses to MarV with responses to all other categories of sounds for each pixel inside the cranial window. Fig. 1d showed the t-value maps and Fig. 1e showed the p-value maps in both marmosets for comparing responses to MarV versus responses to all other categories of sounds. Two restricted regions (referred hereafter to as voice patches) were found in each marmoset that showed significantly stronger responses to MarV than to any other control categories. One patch is located in the posterior core regions and the other one is located in the anterior non-core regions (Fig. 1d-e). To demonstrate the MarV selectivity of a single pixel, responses of two example pixels chosen from marmoset M56E (Fig. 1e, left) are shown in Fig. 1f. The pixel inside the anterior patch showed significantly stronger responses to MarV than other categories of sounds, whereas the pixel outside the anterior patch showed similar responses to all categories of sounds. To further check the robustness of the voice patches, two control analyses were performed. First, t-value maps and p-value maps for comparing responses to MarV versus responses to all other categories of sounds were computed from different numbers of trials in both marmosets (Extended Data Fig. 2). The two voice patches in each marmoset can be computed with a small number of trials, suggesting the good quality of calcium signals and clear MarV selectivity of the voice patches. Second, t-value maps and p-value maps for comparing responses to MarV versus responses to all other categories of sounds were computed from different sound levels in both marmosets (Extended Data Fig. 3). The two voice patches were clearly visible across all sound levels we tested, suggesting the invariance to sound levels.

**Fig. 1.**
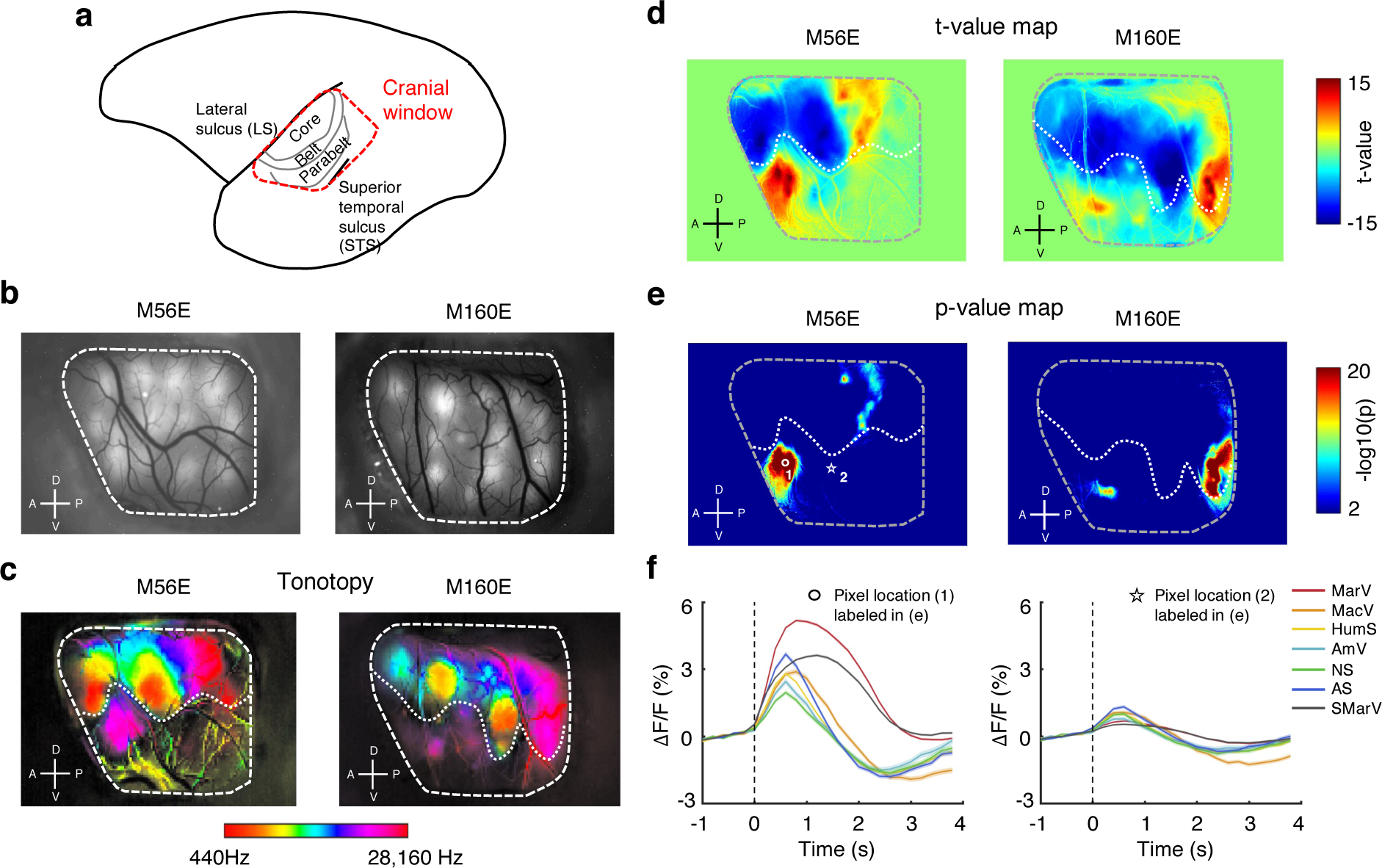
Voice patches in marmoset auditory cortex. **a** An illustration of the location of the implanted cranial window (red dashed contour). The window covers the entire auditory cortex on the brain surface. **b** The implanted cranial windows (white dashed contour) in two marmosets (left hemisphere for M56E and right hemisphere for M160E). Both windows cover the cortical regions between the lateral sulcus and the superior temporal sulcus. The brighter regions inside the windows are the virus injection sites. **c** The tonotopic maps imaged through the cranial windows. The borders between core and non-core regions in the auditory cortex are marked by white dotted lines in both marmosets. **d** The t-value maps in both marmosets derived by comparing the responses to marmoset vocalizations with the responses to all other six categories of sounds. **e** The p-value maps in both marmosets derived by comparing the responses to marmoset vocalizations with the responses to all other six categories of sounds. Two voice patches are found in both marmosets. **f** Response waveforms of two example pixels labeled in (**e**) to all seven categories of sounds (mean ± sem). (MarV: marmoset vocalization; MacV: macaque vocalization; HumS: human speech; AmV: other animal vocalization; NS: natural sounds; AS: artificial sounds; SMarV: scrambled marmoset vocalization).

### Marmoset vocalization selectivity in voice patches

Given the MarV selectivity of the voice patches, we sought to further characterize the response properties of each patch by investigating the vocalization selectivity of all pixels inside the patch. Regions of interest (ROIs) for voice patches were manually drawn based on the p-value maps with p < 0.01 (Fig. 2a for M56E and Extended Data Fig. 4a for M160E). Four ROIs (Posterior Patch 1 (PP-1), Posterior Patch 2 (PP-2), Posterior Patch 3 (PP-3), and Anterior Patch (AP)) were defined in M56E, and three ROIs (Posterior Patch 1 (PP-1), Posterior Patch 2 (PP-2), and Anterior Patch (AP)) were defined in M160E. A Non-Voice Patch (NVP) was defined outside the voice patches with p > 0.05 for each marmoset as a control (Fig. 2a for M56E and Extended Data Fig. 4a for M160E). We then constructed a matrix containing the selectivity profiles of all pixels in each patch. In this matrix, each row represents one pixel, and each column represents one sound stimulus. Figs. 2b, 2e, and 2h show the matrices of normalized intensity change to the baseline intensity (ΔF/F) across all stimuli of all pixels in PP (PP-1, PP-2, and PP-3 were combined together), AP, and NVP respectively in marmoset M56E. Figs. 2c, 2f, and 2i show bar graphs of the average responses to each stimulus across all pixels in PP, AP, and NVP respectively in marmoset M56E. MarV elicited stronger responses across all pixels in PP and AP, but not in NVP, suggesting MarV selectivity in PP and AP. To quantify the MarV selectivity of individual pixels, we computed a MarV selectivity index (MSI, see Methods) for each pixel. MSI measures the distance between responses to MarV and to other sounds. MSI is a value between −1 and 1, it is a positive value if the responses to MarV are higher than the responses to other sounds, and a negative value if the responses to MarV are lower than the responses to other sounds. Positive MSI values were observed in PP and AP, whereas negative values were observed in NVP (Figs. 2d, 2g, and 2j). These results further confirmed the MarV selectivity in PP and AP. Similar results were found in the other marmoset (M160E, Extended Data Fig. 4). Considering multiple sub-regions were included in PP for both marmosets, we next sought to compare the response profiles of all PPs in both marmosets. Extended Data Fig. 5 shows the average responses to each stimulus across all pixels in PP-1 (Extended Data Fig. 5a), PP-2(Extended Data Fig. 5b), and PP-3 (Extended Data Fig. 5c) of marmoset M56E, as well as PP-1 (Extended Data Fig. 5d) and PP-2(Extended Data Fig. 5e) of marmoset M160E. All PP sub-regions in each marmoset showed similar response profiles with the same selectivity to MarV, suggesting similar response patterns of all PP sub-regions. Thus, in the later analysis, all PP sub-regions were combined together.

**Fig. 2.**
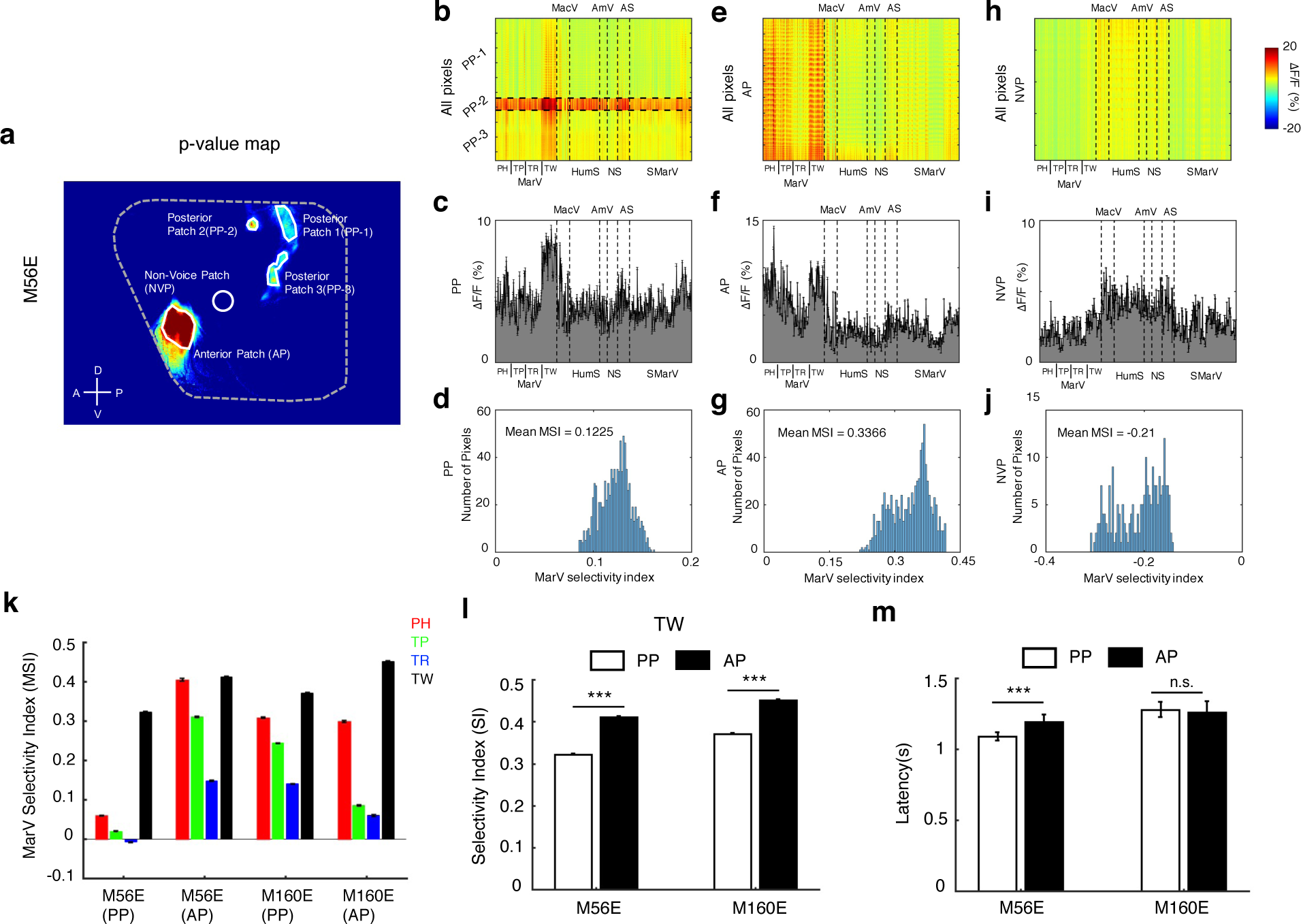
Marmoset vocalization selectivity in voice patches. **a** An illustration of the regions of interest (ROIs, white solid contour) in one marmoset (M56E). ROIs are defined based on the p-value maps. Five ROIs are shown in this marmoset. PP-1: (Posterior Patch 1); PP-2: (Posterior Patch 2); PP-3: (Posterior Patch 3); AP: (Anterior Patch); NVP: (Non-Voice Patch). **b**, **e**, **h** Selectivity profiles of all pixels in all PPs (**b**), AP (**e**), and NVP (**h**) of marmoset M56E to all seven categories of sounds. Each row represents one pixel, and each column represents one sound. To compute selectivity profiles for each pixel, the signal on each pixel is quantified and expressed as the normalized intensity change to the baseline intensity (ΔF/F). **c**, **f**, **i** Average response to each of the seven categories of sounds across all pixels in all PPs (**c**), AP (**f**), and NVP (**i**) of marmoset M56E. Error bars represent ±1 SEM (standard error of mean). **d**, **g**, **j** Distribution of MSIs across all pixels in all PPs (**d**), AP (**g**), and NVP (**j**) of marmoset M56E. **k** MarV selectivity index of four call types in PPs and APs of both marmosets (M56E and M160E). **l** TW selectivity index in PPs and APs of both marmosets (M56E and M160E). **m** Response latencies averaged across all pixels in PPs and APs of both marmosets (M56E and M160E). (***: p < 0.001, rank sum test. PH: Phee; TP: Trillphee; TR: Trill; TW: Twitter).

To check whether the voice patches have different preferences for different call types, MSIs of different call types were calculated for both marmosets. Fig. 2k shows MSI for PH (Phee), TP (Trillphee), TR (Trill), and TW (Twitter) respectively of both marmosets. For both patches in both marmosets, the highest MSI values were found for TW, and the lowest MSI values were found for TR, suggesting the order of MarV call type selectivity from low to high for both patches is TR, TP, PH, and TW. Fig. 2l shows the comparison of TW selectivity for both patches in both marmosets. AP showed significantly higher selectivity index values than those of PP in both marmosets (p < 0.001, rank sum test). We also computed the response latencies of all pixels in both patches. The average response waveforms to all stimuli across all pixels in both patches were shown in Extended Data Fig. 6. Only TW responses were included in the latency analyses since both patches have the highest selectivity to TW. Fig. 2m shows the comparison of peak latencies average across all pixels in both patches of both marmosets. AP showed significantly longer latencies than PP for M56E (p < 0.001, rank sum test), whereas no significant differences were observed for M160E. Together, these results suggest a potential hierarchical organization of PP and AP, in which AP is downstream of PP.

### Call type information is carried by population responses in voice patches

To explore whether voice patches contain information about call types, we examined whether the call type of a marmoset vocalization can be predicted by the population responses of the voice patches. Data from all pixels inside each voice patch with 20 trials were included in this analysis. Template data was derived by randomly picking 10 trials from the original data, the remaining 10 trials were used as validation data. 100 pairs of template data and validation data were constructed by randomly generating the template data for 100 times. Euclidean distance was calculated between template and validation data for each of the stimuli pairs and averaged over 100 times to construct the dissimilarity matrix. Figs. 3a and 3e show the dissimilarity matrix between each of the MarV stimuli in PP and AP for marmoset M56E, respectively. To examine the response similarities for all MarV stimuli, unsupervised multidimensional scaling (MDS) was applied to the dissimilarity matrices to construct a geometric space in which Euclidean distances between different stimuli correspond to the similarities between their neural responses. Figs. 3b and 3f show the MDS results of PP and AP for marmoset M56E, respectively. Visual inspection of the MDS plots suggested that neuronal responses to different marmoset vocalizations are organized by different call types.

**Fig. 3.**
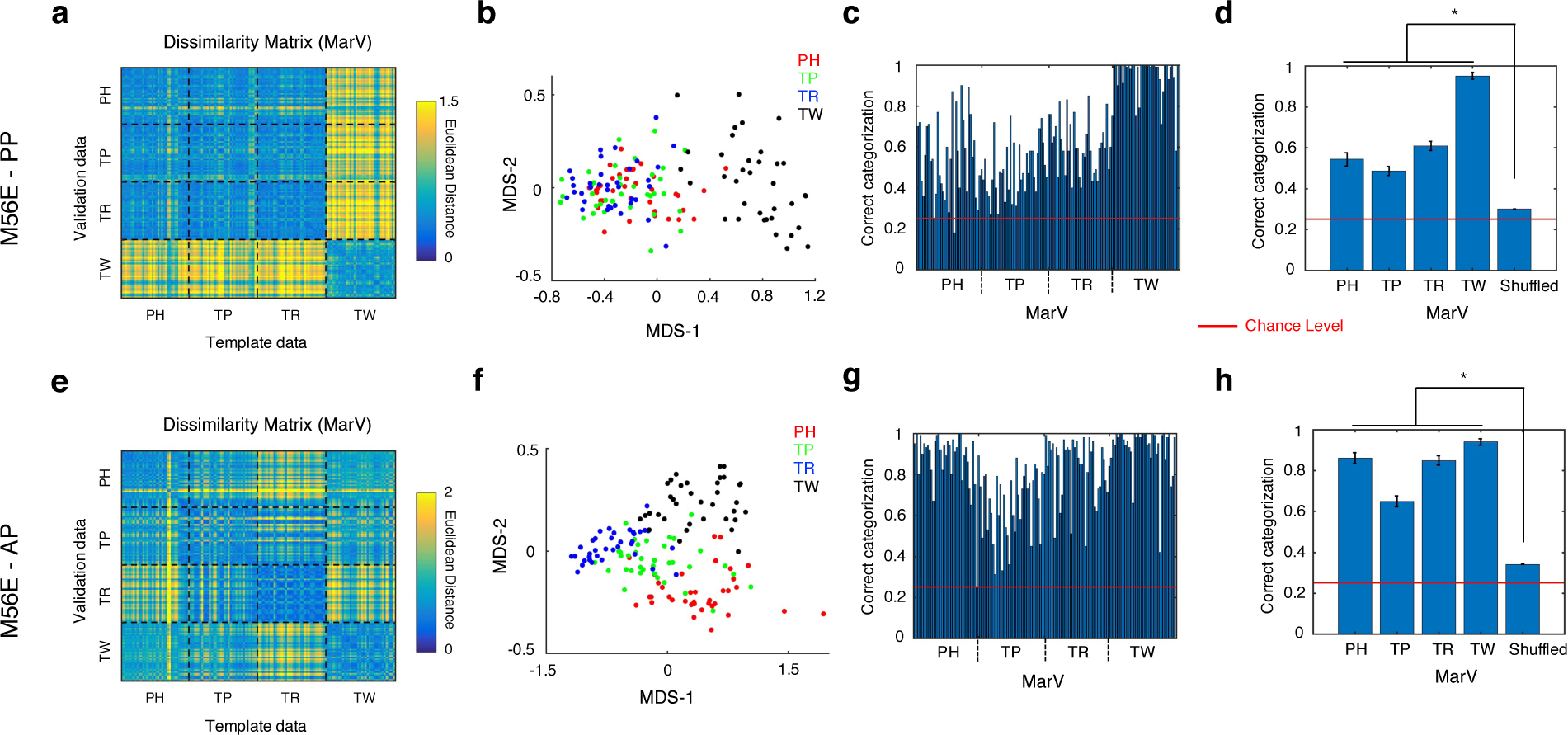
Call types information is carried by population responses in voice patches. **a**, **e** Dissimilarity matrices of Euclidean distances between each of the MarV stimuli pairs of template and validation response data in PP (**a**) and AP (**e**) of marmoset M56E. **b**, **f** Relational organization of response dissimilarity using MDS for call types in PP (**b**) and AP (**f**) of marmoset M56E. **c**, **g** The percentage of correct categorization for call types of each MarV stimulus in PP (**c**) and AP (**g**) of marmoset M56E. **d**, **h** Comparisons of correct categorization between original (grouped by call types) and shuffled data of PP (**d**) and AP (**h**) for marmoset M56E (*: p < 0.05, rank sum test). Chance performance would be 1/4 for call types categorization (indicated by the horizontal red line in each graph).

To confirm the above results, we sought to compute the categorization accuracy of each MarV stimulus based on the template and validation data. If the population responses correctly categorized a particular MarV stimulus as belonging to one of the four call types, then the neuronal response to that MarV stimulus in the validation data should be closest to the response to a MarV stimulus of the same call type in the template data (chance level = 1/4). Figs. 3c and 3g show the percentage of correct categorization for each MarV stimulus of PP and AP for marmoset M56E, respectively. The categorizations of all MarV stimuli are above the chance levels for both PP and AP of marmoset M56E. To validate the robustness of the categorization accuracy, we conducted permutation tests by shuffling labels of the MarV stimuli in validation data for 1000 times. Figs. 3d and 3h show the comparisons of categorization accuracies between the original (grouped by call types) and shuffled data of PP and AP for marmoset M56E, respectively. All stimuli from all four call types in both voice patches can be significantly correctly categorized into categories of call types than those in permutation tests (*: p < 0.05, rank sum test). Similar results were also observed in the other marmoset M160E (Extended Data Fig. 7). Overall, these results suggest that call type information is carried by population responses in voice patches.

### Identity information is carried by population responses in voice patches

Voices and vocalizations carry a wealth of information about the speaker’s identity. To explore whether voice patches contain identity information, we first examined whether discriminant function analysis (DFA) can be utilized to correctly classify marmoset vocalizations to individual-specific groupings based on sets of acoustic features quantitatively measured from those vocalizations. We measured between 9 and 18 acoustic features of the four call types (PH: 9 features; TP: 14 features; TR: 13 features; and TW: 18 features)^33^ for the 144 marmoset vocalizations used in this study. These measured features were physically intuitive parameters, such as dominant frequency, duration, frequency modulation, etc, and were calculated based on the methods published previously^33^. We performed split-sample DFA tests in which 5 of the available 6 vocalizations from each marmoset were used to calculate classification coefficients while the final vocalization was used to test the accuracy of the resulting classifier. Predictor classification was compared with the known identity assignment to determine if it was a correct classification. The DFA tests were performed on every possible subset of the combined acoustic features, and the performance statistics were clustered according to the number of acoustic features in the subset. Fig. 4a shows the DFA performance results as a function of acoustic feature count for the four call types. The highest categorization accuracy is observed for TW, the lowest categorization accuracy is observed for PH, while the categorization accuracies for TP and TR are in between. The categorization accuracies for the four call types are all above chance levels, suggesting that identity information can be significantly decoded based on acoustic features. To determine which specific features provide the most information for discrimination, we extracted the 3 clusters corresponding to the feature sets with the top 3 categorization accuracies for each call type. For each of these clusters, the feature sets comprising the entire cluster were rank ordered according to the discriminant performance, and the top 20 performing feature sets were extracted. A score was then associated with each of these feature sets inversely proportional to its rank order within the cluster (the top-performing feature set was given a score of 20, and the 20^th^-ranked feature set was given a score of 1). Final scores were scaled to contain values ranging from 0 - 1. Extended Data Fig. 8 shows the top 20 performing feature sets from the selected top 3 clusters and the final scores of each acoustic feature for each call type. The most critical acoustic features for discrimination are duration and Max Freq (maximum frequency) for PH (Extended Data Fig. 8d); are duration, End Freq (ending frequency), tMin (time to minimum frequency), and FMdepthMin (minimum difference between a trough and successive peak in a call’s frequency modulation segment) for TP (Extended Data Fig. 8h); are duration, Start Freq (starting frequency), End Freq, tMax (time to maximum frequency), FMdepthMin, and FMrate (frequency modulation rate) for TR (Extended Data Fig. 8l); and are Tphr B (sweep time of beginning phrases), Max Freq M (maximum frequency of middle phrases), Dom Freq M (dominant frequency of middle phrases), Min Freq M (minimum frequency of middle phrases), and Tphr M (sweep time of middle phrases) TW (Extended Data Fig. 8p). These results suggest individual identity can be classified based on sets of acoustic features measured from marmoset vocalizations.

**Fig. 4.**
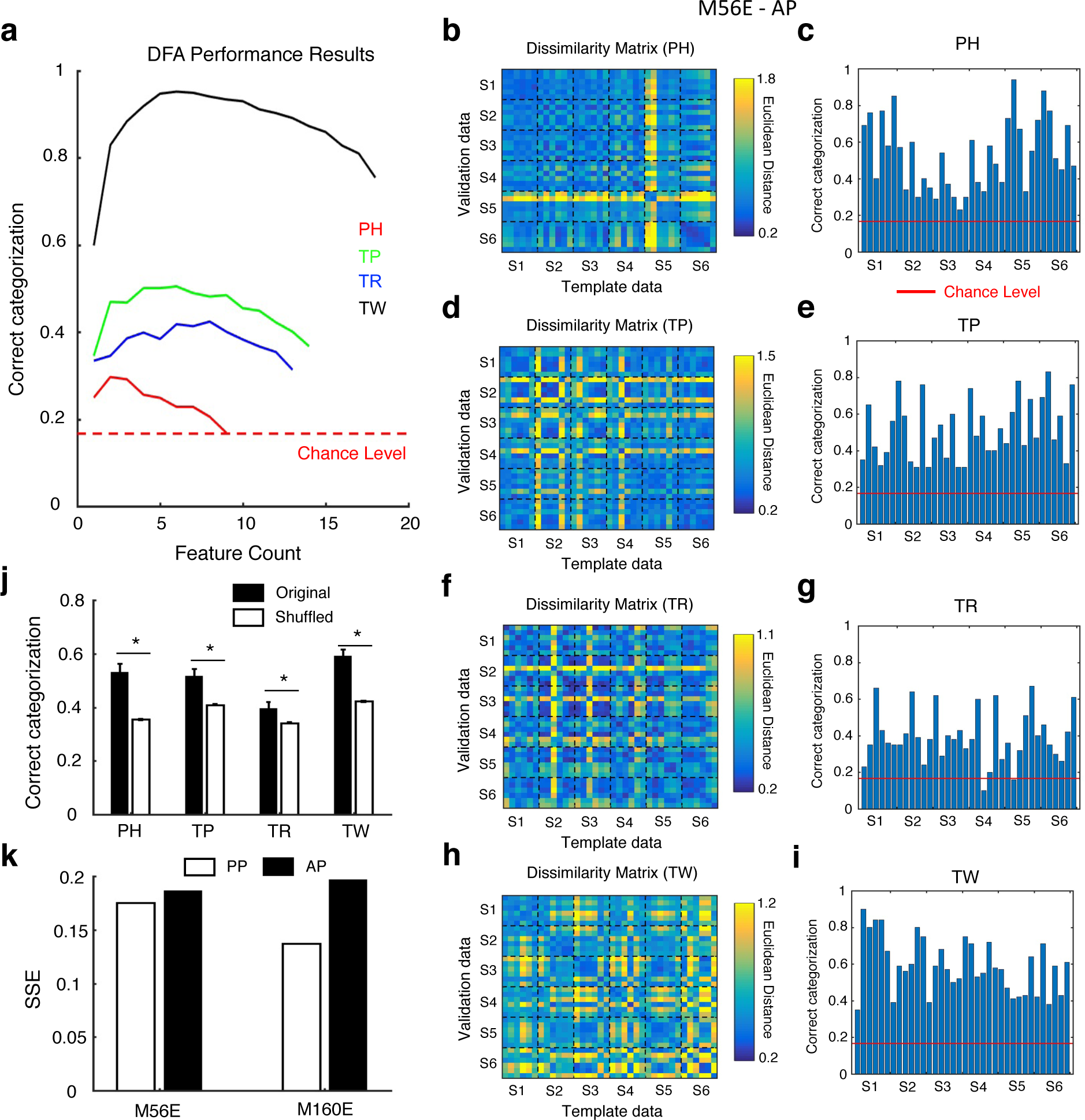
Identity information is carried by population responses in voice patches. **a** DFA performance results as a function of acoustic feature count for the four call types. Chance performance would be 1/6 for identity categorization (indicated by the horizontal red dashed line). **b, d, f, h** Dissimilarity matrices of Euclidean distances between each of the MarV stimuli pairs of test and template response data in AP of marmoset M56E for PH (**b**), TP (**d**), TR (**f**), and TW (**h**). **c, e, g, i** The percentage of correct categorization for identity of each MarV in AP of marmoset M56E for PH (**c**), TP (**e**), TR (**g**), and TW (**i**). **j** Comparisons of correct categorization between original and shuffled data of AP for marmoset M56E (*: p < 0.05, rank sum test). **k** Comparisons of SSE between PP and AP in both marmosets.

We next sought to explore whether identity can be classified based on the neuronal responses in voice patches. We constructed the dissimilarity matrices for each call type using the methods described earlier based on the Euclidean distances calculated between the template and validation data. Figs. 4b, 4d, 4f, and 4h show the dissimilarity matrices between each of the MarV stimuli in AP of marmoset M56E for each call type respectively. Categorization accuracy for each stimulus was then calculated using the same methods described earlier based on the least Euclidean distance (chance level = 1/6 for identity). Figs. 4c, 4e, 4g, and 4i show the percentage of correct categorization for each MarV in AP of marmoset M56E for each call type respectively. All call types can be correctly categorized into categories of identity above the chance levels. To validate the robustness of the categorization accuracy, we conducted permutation tests by shuffling the identity labels of the MarV stimuli in validation data for 1000 times. Fig. 4j shows the comparisons of categorization accuracies between the original and shuffled data of AP for marmoset M56E. All stimuli from all four call types in AP of marmoset M56E can be significantly correctly categorized into categories of identity than those in permutation tests (*: p < 0.05, rank sum test). Similar results were also observed in the PP of marmoset M56E (Extended Data Fig. 9), in the PP of marmoset M160E (Extended Data Fig. 10), and in the AP of marmoset M160E (Extended Data Fig. 11). Overall, these results suggest that identity information is carried by population responses in voice patches.

To quantify the distance between neuronal categorization and acoustic categorization for individual identity, we calculated SSE (Sum of Squared Error, see Methods) across the four call types. A higher SSE value indicates a larger distance between neuronal and acoustic categorizations. Fig. 4k shows the comparisons of SSE values between PP and AP in both marmosets. PP has a smaller SSE value than that of AP in both marmosets, suggesting neuronal categorization is closer to acoustic categorization in PP than in AP. These results agree with our findings that PP and AP are hierarchically organized, in which AP is downstream of PP.

### Connectivity of voice patches

In the visual system, face patches are interconnected and form a specifically hierarchical network^34^. To check whether voice patches have similar properties, we sought to investigate the connectivity of voice patches. We compared the similarity of responses of each voice patch using the co-activation method^6, 35^ by correlating the maximum amplitudes across all trials under the TW condition (All voice patches showed the highest selectivity to TW). Fig. 5a shows the positions of the four example pixels chosen from M56E. Pixel 1 is located in the AP and is chosen as the example reference pixel. Pixel 2 is located in the AP and is within the same voice patch as Pixel 1 (within-patch pair). Pixel 3 is located in the PP and is in a voice patch different from Pixel 1 (between-patch pair). Pixel 4 is located in the NVP and is outside of any voice patches (outside-patch pair). Fig. 5b shows the maximum response amplitudes across all trials under TW condition for the four example pixels. Pearson’s correlations between the reference Pixel 1 and the other three pixels were then calculated (Fig. 5c), and the significance was confirmed by permutation tests (P < 0.05 was chosen as the criterion). We observed significant correlations between all pixel pairs (1-2: within-patch pair; 1-3: between-patch pair; 1-4: outside-patch pair). Moreover, the correlation between the within-patch pair was the highest while the correlation between the outside-patch pair was the lowest. To generalize this observation, the correlation analysis was applied to all pixel pairs for each marmoset. By averaging correlations across all within-patch pixel pairs, all between-patch pixel pairs, and all outside-patch pixel pairs for each marmoset, we observed that the correlations for within-patch and between-patch pixel pairs were significantly higher than those for outside-patch pixel pairs and that the correlations for within-patch pixel pairs were significantly higher than those for between-patch pixel pairs in both marmosets (Fig. 5d, P < 0.001, rank sum test). Taken together, these results indicate that pixels within a voice patch and between voice patches are highly connected, suggesting similar cortical architectures of face patches and voice patches.

**Fig. 5.**
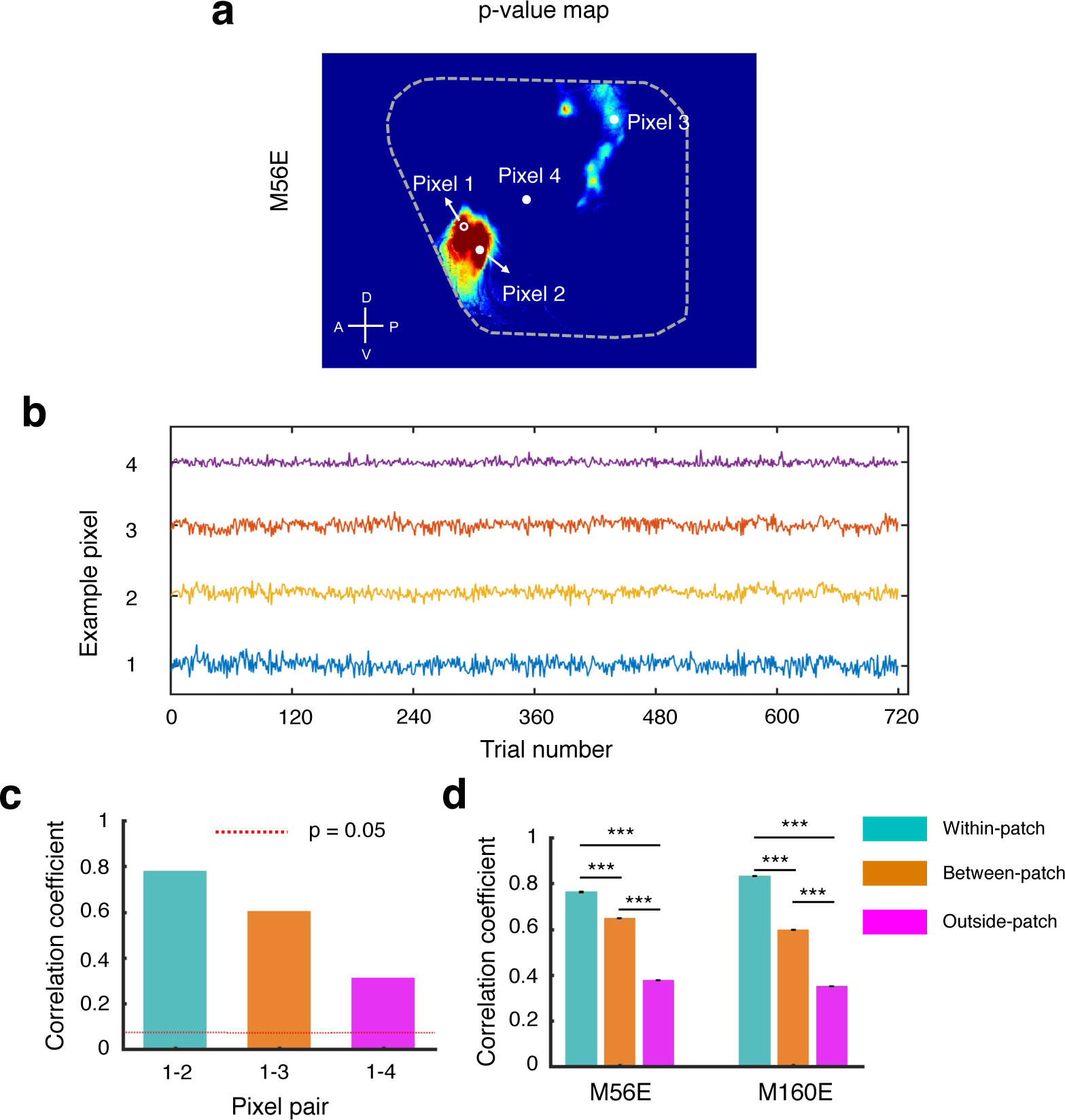
Connectivity of voice patches. **a** Four example pixels are chosen (Pixel 1: reference pixel in AP; Pixel 2: within-patch pixel of Pixel 1; Pixel 3: between-patch pixel of Pixel 1; Pixel 4: outside-patch pixel of Pixel 1). **b** Maximum responses of the four example pixels across all trials under TW condition. **c** Correlations between the reference pixel (Pixel 1) and the other three pixels (significance levels are determined by permutation tests). **d** Comparisons of correlations from all within-patch pixel pairs, all between-patch pixel pairs, and all outside-patch pixel pairs in both marmosets (***: p < 0.001, rank sum test).

## DISCUSSION

### The existence of voice patches for processing species-specific vocalizations in marmosets

In the visual system, face patches were found to be specialized for processing faces in humans and non-human primates^13–15^. The face patches consist of a series of discrete and inter-connected cortical regions that are selective to faces over other images. Using wide-field calcium imaging, we demonstrated the existence of voice patches that prefer marmoset vocalizations over other sounds in the marmoset auditory cortex. We found two voice patches with one located in the posterior core regions and the other one located in the anterior non-core regions (Fig. 1). These voice patches are hierarchically organized (Fig. 2) and functionally connected (Fig. 5). Moreover, information about call type and identity is found to be available within the voice patches (Figs. 3, 4). These results provide insights into understanding how species-specific vocalizations are processed in the marmoset brain, suggesting similar cortical architectures for processing faces and vocalizations.

Previous studies have demonstrated the existence of vocalization-specific regions in macaques^16^ and marmosets^17^ using fMRI. One concentrated vocalization-specific region was found to be located in the anterior temporal lobe outside the auditory cortex for each hemisphere in both macaques and marmosets. Our findings of multiple voice patches inside the auditory cortex suggest a discrepancy with the previous studies. The following reasons may explain this discrepancy. Firstly, previous studies used fMRI to study vocalization processing in non-human primates. The substantial acoustic noises generated by fMRI sequences^18^ have profound impacts on auditory experiments. The echo-planar imaging (EPI) sequence used in these fMRI experiments can generate sound pressures as high as 110-130 dB^36–37^, which can largely mask the neuronal responses to vocalizations presented at 60 dB SPL. Even though sparse sampling imaging technique^19^ has been widely used in auditory neuroscience research^16–17^, it is at a cost of reduced temporal resolution, resulting in fewer repetitions with a lower signal-to-noise ratio (SNR). Thus, the fMRI signals may not have adequate SNR to locate the voice patches in the auditory cortex. Secondly, fMRI has a relatively low spatial resolution compared with our wide-field calcium imaging. The maximum size of the voice patches found in this study is around 1.5×1.5 mm. The in-plane spatial resolution of the fMRI study on marmosets^17^ is 0.625×0.625 mm, which means only 6 pixels will be captured by the fMRI for one of the largest voice patches found in the current study. The auditory cortical regions in macaques have expanded between 8X to 16X in three dimensions compared with those in marmosets^38^, suggesting estimates of 3×3 mm to 3.8×3.8 mm for the largest voice patches found in the current study. Considering the in-plane spatial resolution of the fMRI study on macaques^16^ is 1×1 mm, only 9 to 14 pixels will be detected by the fMRI for one of the largest voice patches found in the current study in macaques. These small expected number of pixels might thus make the specialized cortical regions difficult to detect. Lastly, the implanted cranial window used in the current study only covers the auditory cortex including core, belt, and parabelt regions^39^. The vocalization-specific regions in macaques^16^ and marmosets^17, 40^ found previously using fMRI were located in the anterior temporal lobe, which is outside the regions within the cranial window. We expect that there is another voice patch in the anterior temporal lobe. Future studies with cranial windows implanted in this region will help to clarify this question.

### Evolution of the voice patches

In non-human primates, the auditory information is thought to be processed through two parallel hierarchical streams, including a ventral “what” stream and a dorsal “where” stream^39^. In humans, two parallel hierarchical streams are also proposed for speech processing, it contains a dorsal pathway for auditory-motor integration, which is different from the dorsal “where” stream in non-human primates^41–43^. In humans, three voice patches (posterior, middle, and anterior voice patch) located in the superior temporal gyrus (STG) were found both using fMRI^5^ and electrocorticographic (ECoG)^6^. These three voice patches are hierarchically organized through a dual-directional information flow, in which the middle voice patch functions as the initial hub and anterior and posterior voice patches are downstream of the middle voice patch^6^. Our results also suggest a hierarchical organization of the voice patches in marmosets. We observed an increase in MarV selectivity (Fig. 2l) and an increase in response latencies (Fig. 2m) from the posterior to anterior voice patch. These results suggest homologous brain structural and functional organizations for processing species-specific vocalizations in marmosets and humans.

However, a comparison of the anatomic locations of the voice patches between marmosets and humans suggests two discrepancies. Firstly, a voice patch was found to be located in the core regions of marmosets, while no voice patch was found in the primary auditory cortex of humans. Secondly, three voice patches per hemisphere were found in humans while only two voice patches were found in marmosets. The possible explanations are as follows. Firstly, the primary auditory cortex in humans is located inside the lateral sulcus^44–46^. ECoG electrodes cannot reach that area and the fMRI signal has a bad SNR in that area, which thus makes the possible voice patch in the primary auditory cortex hard to detect. Secondly, the implanted cranial window used in the current study only covers the auditory cortex. We highly expect two more voice patches located in the anterior and posterior temporal regions respectively. Overall, if this is the case, the temporal lobes of both humans and marmosets will have four voice patches. One is located in the primary auditory cortex, the other three are located in high-level auditory regions. These voice patches are likely to be hierarchically organized through a dual-directional information flow, in which the voice patch in the primary auditory cortex serves as the first stage, the middle voice patch in the high-level auditory regions serves as the inter-medial stage, and the anterior and posterior voice patches in the high-level auditory regions serve as the last stages. Alternatively, the voice patches in the two primates were repositioned following the lineages’ split from their common ancestor. Further comparative studies of structure and function in multiple primate species will be critical to understanding the evolution of voice patches in primates.

### Hemispherical asymmetries of the voice patches

Hemispherical asymmetries of the auditory cortex have been proposed based on anatomic studies^47–48^, which showed the asymmetries of the afferent pathways. The anatomic asymmetries lay the foundations for the functional asymmetries of the auditory cortex. Substantial evidence exists indicating that, in humans, speech sounds are preferentially processed in the left auditory regions, while music sounds are processed in the right auditory regions. In addition, the left auditory regions process temporal information, while the right auditory regions process spectral information^47–50^. The previous study on voice patches in humans has demonstrated the existence of left lateralization in processing voices^6^. The lateralization is mainly focused on two aspects. First, left lateralization of native language processing. Second, left lateralization of auditory-motor interaction in speech perception. These studies suggest that voice patches in both hemispheres may be dedicated to processing different information buried in the voices.

In the current study, we studied the voice patches in two marmosets with cranial windows implanted in different hemispheres. Cranial windows were implanted in the left hemisphere for M56E and in the right hemisphere for M160E. We observed substantial differences between the voice patches found in these two marmosets (Fig. 1). For M56E, the anterior voice patch showed significantly better selectivity to MarV than the posterior voice patch, whereas, M160E showed opposite results (Fig. 1e). These results may indicate hemispherical asymmetries of the voice patches. Since the anterior voice patch is downstream of the posterior voice patch, better MarV selectivity in the anterior voice patch of the left hemisphere may suggest left lateralization of higher-level vocalization processing. However, we cannot conclude the hemispherical asymmetries of the voice patches since only two marmosets were included in this study. In addition, the virus was injected into the cortex through several injection sites. The diffusion of the virus in the cortex may lead to inhomogeneous expressions of GCaMP6, which may result in different response amplitudes in different regions. Further studies using transgenetic marmosets with a sufficient sample size can help to clearly elucidate this question.

## METHODS

### Marmoset preparation and wide-field calcium imaging setups

All experimental procedures and methods were approved by the Johns Hopkins University Animal Use and Care Committee and conformed to local and US National Institutes of Health guidelines. Two adult (both females, M56E and M160E) marmoset monkeys (*Callithrix jacchus*) were included in the current study. Three procedures were performed for each marmoset. Firstly, chronic head-cap implantation. The head-cap design and implantation surgery were previously described^51^. During this procedure, two stainless steel head posts were attached to the top of the skull with dental cement for head fixation, and two recording chambers atop each side of the auditory cortex were built. Secondly, cranial window implantation. The artificial dura-based cranial window implantation procedure was previously described^28–29, 52^. The artificial dura is made of silicone and is removable for maintenance under anesthesia. The artificial dura was implanted to cover the entire auditory cortex. The gap between the craniotomy edge and the sidewall of the artificial dura was filled and sealed with surgical silicone adhesive (WPI, Kwik-Sil). M56E has a cranial window implanted in the left hemisphere, and M160E has a cranial window implanted in the right hemisphere. Lastly, virus injections. The cortex-based virus injection procedure was previously described^29^. We injected AAV-DJ-CamKII-GCaMP6 (UNC vector core) into the cortex. The CamKII promoter was reported to control the expressions of cortical pyramidal neurons^53^. In vivo calcium imaging was conducted at least 3 weeks after virus injections.

Wide-field calcium imaging was performed using the XINTRINSIC setups developed in our lab. The details for XINTRINSIC setups were previously described^28–29, 52^. Recordings were acquired using custom code in MATLAB (Mathworks, https://github.com/x-song-x/XINTRINSIC), with a resolution of 1920 × 1080 and a sampling rate of 20Hz (20 frames per second). The marmosets were trained to sit in a custom-designed marmoset chair following the adaptation protocol previously described^51^. Both marmosets were head-fixed and awake during the imaging sessions.

### Stimuli and experimental design

Stimuli consist of seven categories of sounds. 1) marmoset vocalization (MarV): the MarVs were recorded from six marmosets from the colony. Four types of calls (PH, TP, TR, and TW) were included for each marmoset. Each call type for each marmoset has six samples. In total 144 MarVs (6 × 4 × 6) were included. 2) macaque vocalization (MacV): The MacVs were chosen from the Rhesus Monkey Repertoire provided by Dr. Marc Hauser at Harvard University^54^. Ten major call types (Coo, Aggressive, Gecker, Scream, Girney, Shrill bark, Warble, Harmonic arch, Copulation scream, Grunt) were included with each call type having three samples. In total 30 MacVs (10 × 3) were included. 3) Human speech (HumS, 70 stimuli). 4) Other animal vocalization (AmV, 18 stimuli). 5) Natural sounds (NS, 24 stimuli). 6) Artificial sounds (AS, 28 stimuli), and 7) scrambled marmoset vocalization (SMarV): SMarVs are the phase scramble of the MarVs to preserve the spectral content (144 stimuli). HumS, AmV, NS, and AS stimuli were derived from the “Animal, Artificial, Natural, Speech, and Vocal Non-Speech sounds” dataset^55^. All stimuli had roughly similar durations (1.091 ± 0.593 s). All stimuli were normalized to the same amplitude level (60 dB SPL). To test the effects of sound levels on voice patches, we also presented the stimuli to the marmosets at 40 dB SPL and 80 dB SPL. The presentation of the normalized stimuli was controlled by XINTRINSIC in MATLAB (MathWorks). The stimuli (16 bits, 100,000 sampling rate) were attenuated by a programmable attenuator (Tucker-Davis Technology, PA5) and were delivered via a loudspeaker (KEF, LS50) placed ∼1 m away in front of the marmoset. All stimuli were presented in a randomized order with each repeated 20 times. The length of each trial was 5 seconds with a 1 second baseline. The marmosets stayed awake with head-fixed inside a double-walled soundproof chamber (Industrial Acoustics, custom model) during the imaging sessions.

### Calcium imaging data preprocessing and statistic maps

All analyses were performed using MATLAB (MathWorks). The calcium imaging data was first downsampled to a resolution of 480 × 270 and a sampling rate of 5Hz (5 frames per second). Motion corrections were then performed using NoRMCorre^56^ (rigid registration). The first frame of each session was used as a template for registration. The calcium signal for each pixel was quantified and expressed as a normalized fluorescence change to a baseline fluorescence value *(ΔF/F = (F – F0)/F0*). *F0* is the mean fluorescence value during the baseline, and *F* is the raw fluorescence value. The average of *ΔF/F* across trials for each stimulus was taken as a measurement of response amplitude to that stimulus.

t-value maps and p-value maps were derived by comparing the response amplitudes to marmoset vocalizations with the responses to all other six categories of sounds. Welch’s t-test was used to generate the t-value and the p-value for each pixel. t-value maps and p-value maps were shown by combining the t-statistic of all pixels.

### Marmoset vocalization selectivity index and latency

To quantify the MarV selectivity of each pixel, we defined a MarV selectivity index (MSI) for each pixel as the following:

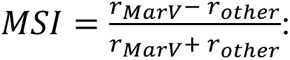

*r_marV_*: mean response amplitude to MarV
*r_other_*: mean response amplitude to all other categories of sounds

MSI measures the distance between responses to MarV and to other sounds. MSI is a value between −1 and 1, it is a positive value if the responses to MarV are higher than the responses to other sounds, and a negative value if the responses to MarV are lower than the responses to other sounds.

MarV response latency was measured after the stimulus onset using the *ΔF/F* waveform of each trial. Latency was defined as the time point at the peak of the response waveform. Only TW responses were included in the latency analyses since both patches have the highest selectivity to TW.

### Categorization of identities from marmoset vocalizations based on acoustic features

Discriminant function analysis (DFA) utilizes multiple independent variables to predict group membership according to specific, categorical, dependent variables. We used linear discriminant functions to test whether we could generate a discriminant model to correctly classify natural vocalizations to individual-specific groupings based on sets of acoustic features quantitatively measured from those vocalizations.

We measured between 9 and 18 acoustic features of the four most commonly produced marmoset call types^33^ for all 144 marmoset vocalizations used in this study. Specifically, 9 features including duration, Max Freq (maximum frequency), Min Freq (minimum frequency), Dom Freq (dominant frequency), Start Freq (starting frequency), End Freq (ending frequency), tMax (time to maximum frequency), tMin (time to minimum frequency), and BW (bandwidth) were calculated for PH. 14 features including duration, Max Freq, Min Freq, Dom Freq, Start Freq, End Freq, tMax, tMin, BW, FMdepthMax (maximum difference between a trough and successive peak in a call’s frequency modulation segment), FMdepthMin (minimum difference between a trough and successive peak in a call’s frequency modulation segment), FMdepth (frequency modulation depth), FMrate (frequency modulation rate), and tTrans (time of transition from sinusoidal to linear FM) were calculated for TP. 13 features including duration, Max Freq, Min Freq, Dom Freq, Start Freq, End Freq, tMax, tMin, BW, FMdepthMax, FMdepthMin, FMdepth, and FMrate were calculated for TR. 18 features including duration, Nphr (number of discernible voicing segments), IPI (inter-phrase interval), Max Freq B (maximum frequency of beginning phrases), Min Freq B (minimum frequency of beginning phrases), Dom Freq B (dominant frequency of beginning phrases), BW B (bandwidth of beginning phrases), Tphr B (sweep time of beginning phrases), Max Freq M (maximum frequency of middle phrases), Min Freq M (minimum frequency of middle phrases), Dom Freq M (dominant frequency of middle phrases), BW M (bandwidth of middle phrases), Tphr M (sweep time of middle phrases), Max Freq E (maximum frequency of ending phrases), Min Freq E (minimum frequency of ending phrases), Dom Freq E (dominant frequency of ending phrases), BW E (bandwidth of ending phrases), and Tphr E (sweep time of ending phrases) were calculated for TW. These acoustic features were calculated based on the methods published previously^33^. Acoustic feature data were then linearly scaled between 0 – 1 to ensure all values were on the same order of magnitude.

We performed split-sample DFA tests in which 5 of the available 6 vocalizations from each marmoset were used to calculate classification coefficients while the final vocalization was used to test the accuracy of the resulting classifier. Predictor classification was compared with the known identity assignment to determine if it was a correct classification. We conducted this analysis 50 times for each set of vocalizations. Once all predictor data had been processed, the averaged percentage of correct classifications from the discriminant function results was calculated as a performance metric.

To determine which subset of features provides the most information in making these distinctions, we tested every possible subset of the combined scaled features and clustered performance statistics according to the number of features in the subset (i.e., performance results for all feature subsets consisting of two features were in cluster two, performance results for all feature subsets consisting of three features were in cluster three). The best performance statistic in each cluster was extracted and compared with other clusters’ best performance statistics. We defined the feature subset with the best performance as the optimum feature set size. We assumed that this set of features thus conveys most of the information required to adequately discriminate identities.

To determine which specific features provide the most information for discrimination, we extracted the 3 clusters corresponding to the feature sets with the top 3 categorization accuracies for each call type. For each of these clusters, the feature sets comprising the entire cluster were rank ordered according to the discriminant performance, and the top 20 performing feature sets were extracted. A score was then associated with each of these feature sets inversely proportional to its rank order within the cluster (the top-performing feature set was given a score of 20, and the 20^th^-ranked feature set was given a score of 1). These scores were attributed to the specific features comprising the feature sets and a cumulative score was calculated for each feature by summing the scores for that feature across all three clusters. With this scoring system, the highest score achievable is 630. Final scores were scaled to contain values ranging from 0 - 1. We defined notable features for each call type as those with a cumulative score of at least 0.5 (i.e., half the maximum possible score) and critical features as those with scores of at least 0.75.

### Calculation of neuronal response dissimilarity

We estimated the dissimilarity of the neuronal response patterns evoked by different stimuli using the Euclidean distances across all pixels inside a voice patch between the template and validation data. Template data was derived by randomly picking 10 trials from the original data of 20 trials, the remaining 10 trials were used as validation data. 100 pairs of template data and validation data were constructed by randomly generating the template data for 100 times. Euclidean distance was calculated between the template and validation data for each of the stimuli pairs and averaged over the 100 times to construct the dissimilarity matrix.

Unsupervised multidimensional scaling (MDS) was applied to the dissimilarity matrices to represent neuronal responses to different MarV stimuli in a new space in which the distance between neuronal responses represents their relative dissimilarity^57^.

### Categorization of marmoset vocalizations based on neuronal responses

The categorization of MarVs based on neuronal responses was performed in two conditions: 1) Group into call type categories (4 call types); 2) Group into identity categories (6 identities). If the population responses in the voice patch correctly categorize a MarV stimulus as belonging to one of the categories (call type or identity), the neuronal response to that MarV stimulus in the validation data should be closest to the response to a MarV stimulus of the same category in the template data (chance level = 1/4 for call type, chance level = 1/6 for identity). Permutation tests were conducted by shuffling labels of the MarV stimuli in validation data for 1000 times.

### Calculation of Sum of Squared Error

SSE (Sum of Squared Error) was calculated as:

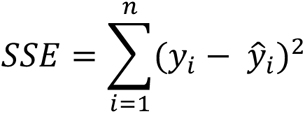

Where:

*n_i_*: number of call types

*y_i_*: neuronal categorization accuracy for call type *i*

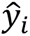: acoustic categorization accuracy for call type *i*

SSE measures the distance between neuronal categorization and acoustic categorization across the four call types. A higher SSE value indicates a larger distance.

### Connectivity of voice patches

The connectivity of voice patches was performed using the co-activation method^6, 35^. We compared the similarity of responses of each voice patch by correlating the maximum amplitudes across all trials under the TW condition (all voice patches showed the highest selectivity to TW). For all data, correlation values were estimated via Pearson’s correlation coefficient. The significance level was computed by shuffling the order of the trials 1000 times. Rank sum test was used to compare the correlation values across different voice patch pairs.

## ACKNOWLEDGEMENTS

This research was supported by National Institute of Health grant DC003180 (X.W.). We thank J. Lynch, K. Schonvisky, S. Miller, E. Easter, and J. Izzi for their assistance with surgeries and animal care, Linyun Zhao for sharing the acoustic feature analysis code, and all Wang’s lab members for their comments on the manuscript.

## AUTHOR CONTRIBUTIONS

Y.Z. and X.W. designed the study. X.S., Y.G., C.C., and X.W. developed the marmoset optical imaging preparation. X.S. developed the XINTRINSIC setup. M.S.O. performed the acoustic discriminant function analysis. Y.Z. performed the experiments and analyzed data. Y.Z. and X.W. interpreted data and wrote the manuscript with comments from all co-authors.

## DATA AVAILABILITY

The raw data supporting the current study have not been deposited in a public repository because of the large size of the dataset but are available from the corresponding authors upon reasonable request.

