## Supplemental Documents for "Voice patches in the marmoset auditory cortex revealed by wide-field calcium imaging"

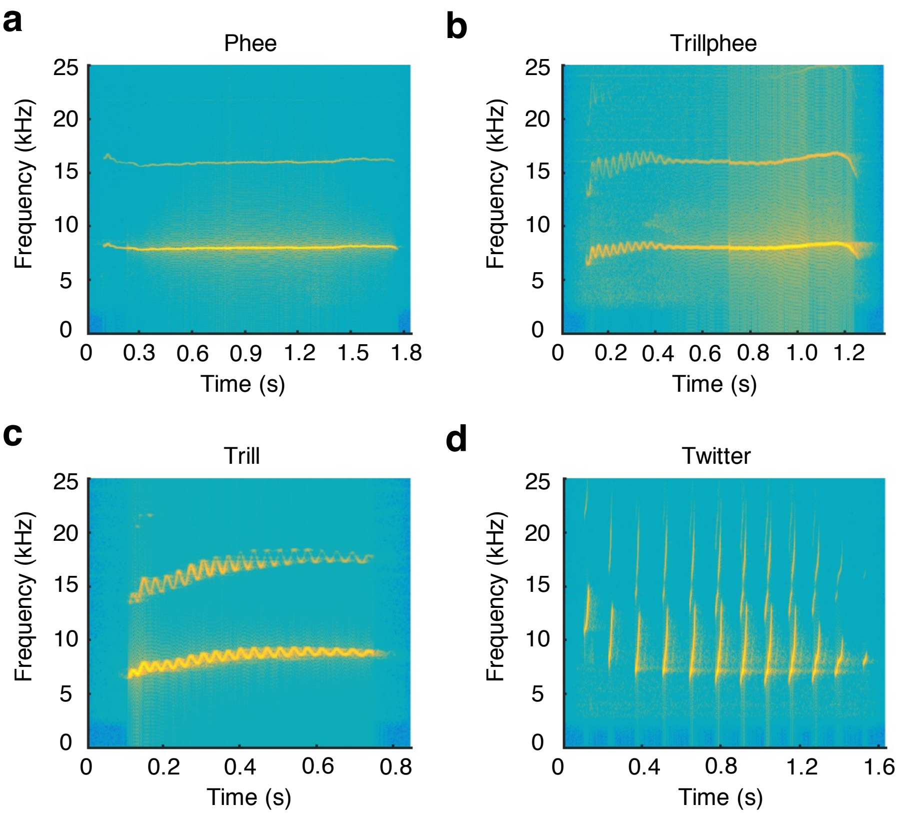


**Extended Data Fig. 1 Example spectrograms of the four most commonly produced marmoset vocalizations. a** Phee. **b** Trillphee. **c** Trill. **d** Twitter.

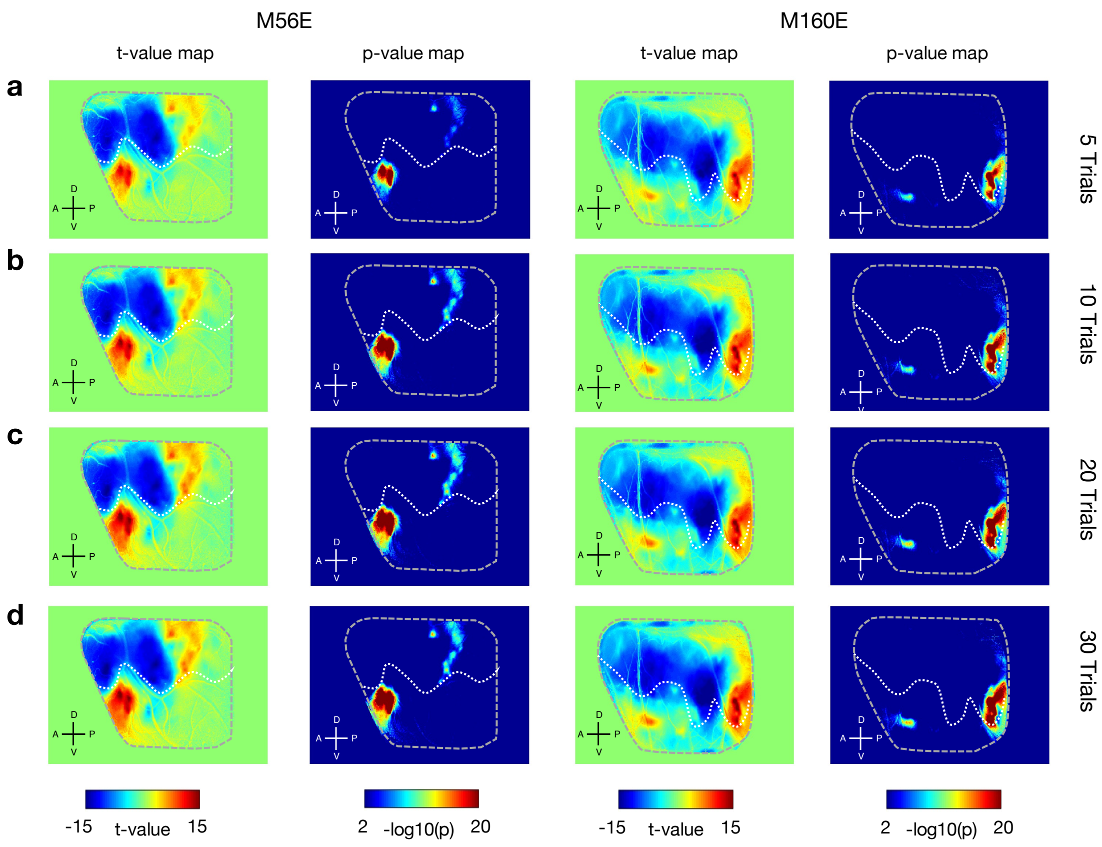


**Extended Data Fig. 2 Voice patches can be computed with a small number of trials.** t-value maps and p-value maps are computed from five trials (**a**), ten trials (**b**), twenty trials (**c**), and thirty trials (**d**) in both marmosets.

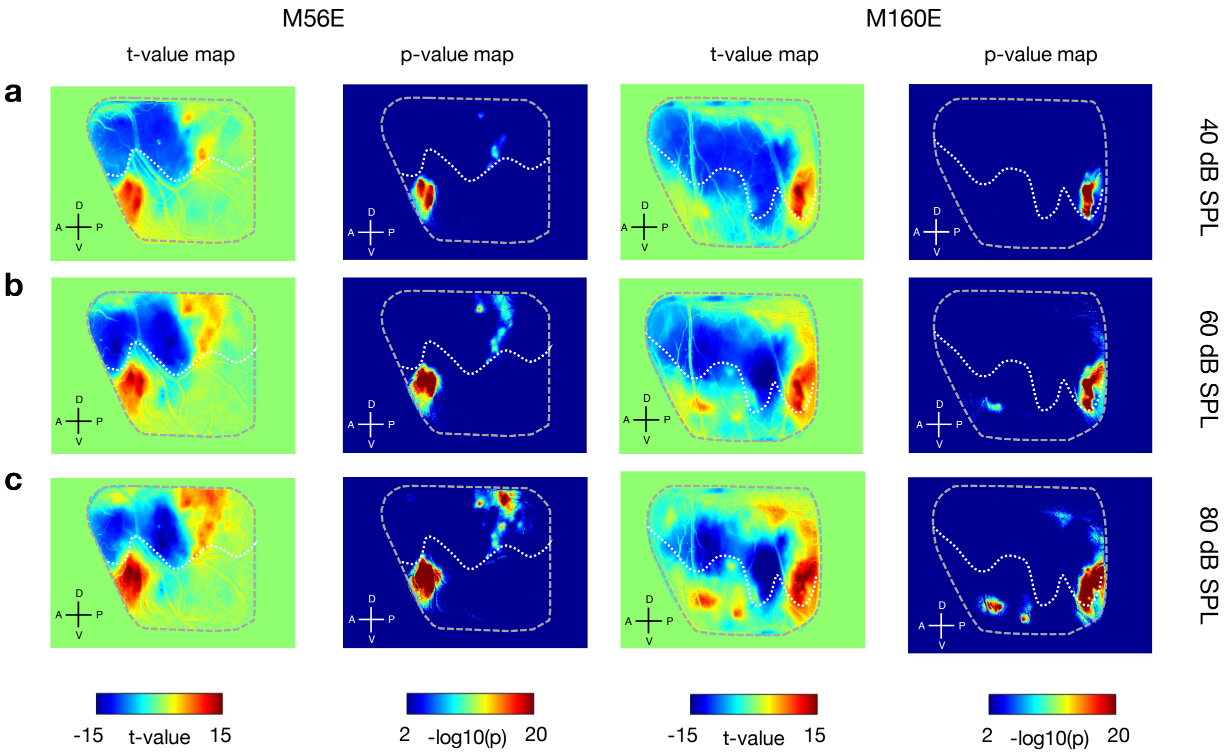


**Extended Data Fig. 3 Voice patches are invariant to sound levels.** t-value maps and p-value maps are computed with the sound levels of 40 dB SPL (**a**), 60 dB SPL (**b**), and 80 dB SPL (**c**) in both marmosets.

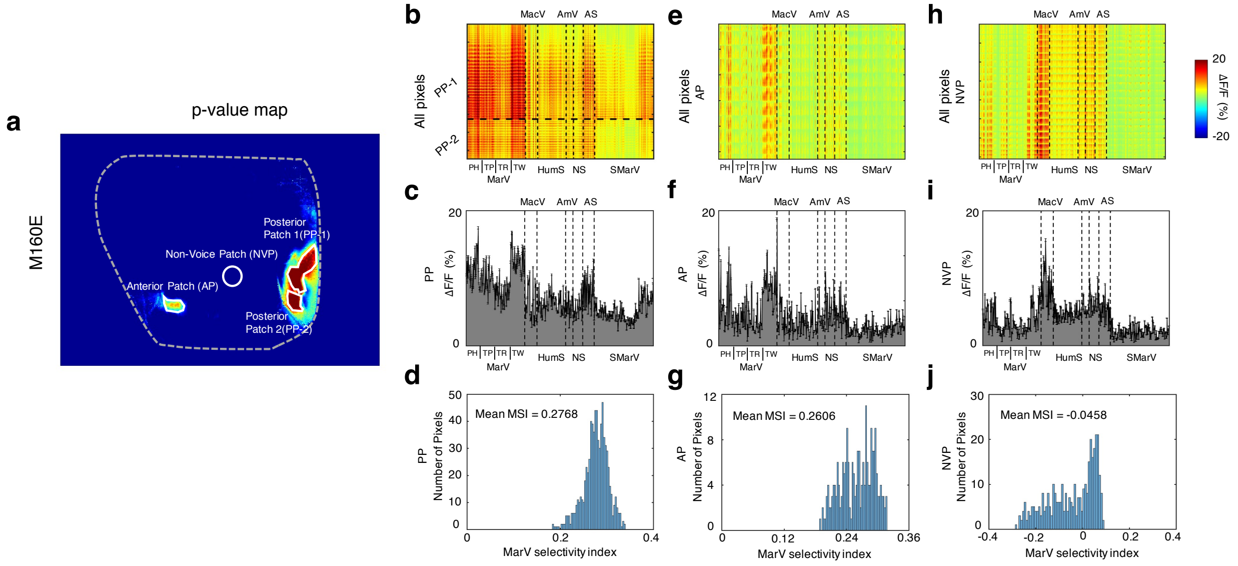


### Extended Data Fig. 4 Marmoset vocalization selectivity in voice patches of marmoset M160E.

**a** An illustration of the regions of interest (ROIs, white contour) in marmoset M160E. **b**, **e**, **h** Selectivity profiles of all pixels in all PPs (**b**), AP (**e**), and NVP (**h**) of marmoset M160E to all seven categories of sounds. **c**, **f**, **i** Average response to each of the seven categories of sounds across all pixels in all PPs (**c**), AP (**f**), and NVP (**i**) of marmoset M160E. **d**, **g**, **j** Distribution of MSIs across all pixels in all PPs (**d**), AP (**g**), and NVP (**j**) of marmoset M160E.

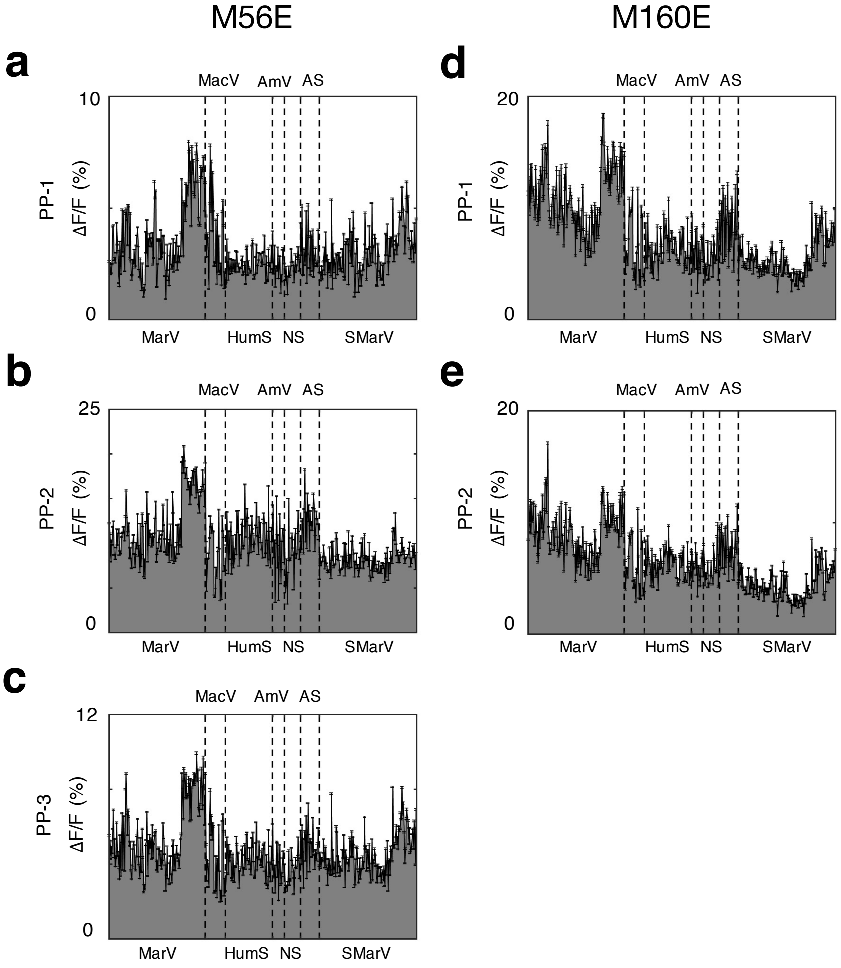


**Extended Data Fig. 5 Response profiles of all PPs in both marmosets.**

**a**, **b**, **c** Average response to each of the seven categories of sounds across all pixels in PP-1 (**a**), PP-2 (**b**), and PP-3 (**c**) of marmoset M56E. **d**, **e** Average response to each of the seven categories of sounds across all pixels in PP-1 (**d**) and PP-2 (**e**) of marmoset M160E. Error bars represent ±1 SEM.

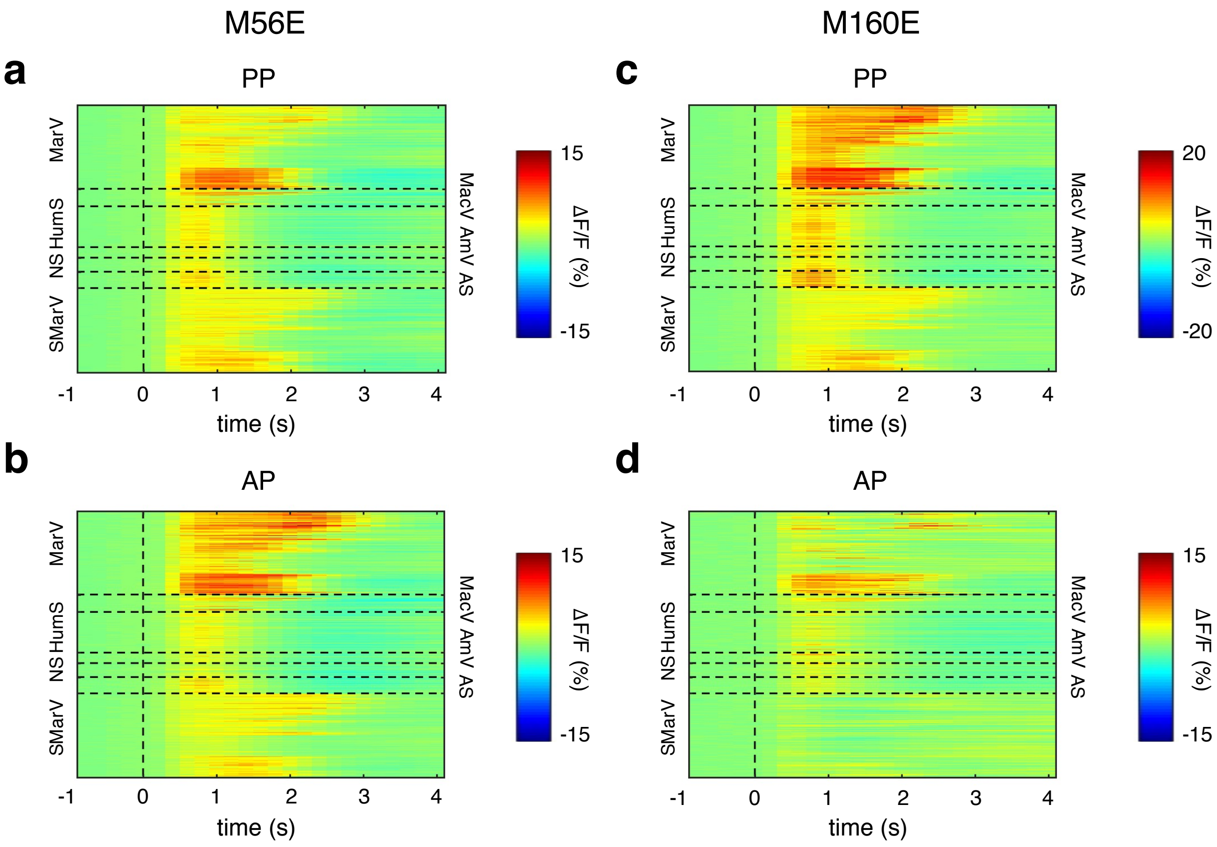


**Extended Data Fig. 6 Average response waveforms in voice patches.**

**a**, **b** Average response waveforms to each of the seven categories of sounds across all pixels in PP (**a**) and AP (**b**) of marmoset M56E. **c**, **d** Average response waveforms to each of the seven categories of sounds across all pixels in PP (**c**) and AP (**d**) of marmoset M160E.

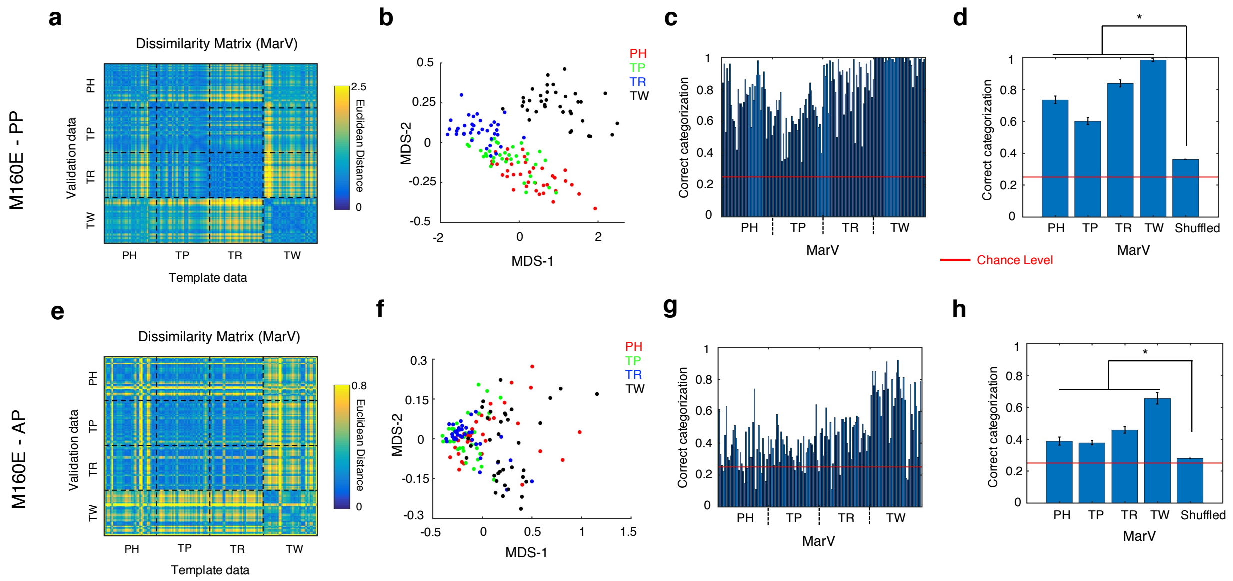


**Extended Data Fig. 7 Similar results are computed in marmoset M160E as the results in marmoset M56E shown in Fig. 3.**

**a**, **e** Dissimilarity matrices of Euclidean distances between each of the MarV stimuli pairs of template and validation response data in PP (**a**) and AP (**e**) of marmoset M160E. **b**, **f** Relational organization of response dissimilarity using MDS for call types in PP (**b**) and AP (**f**) of marmoset M160E. **c**, **g** The percentage of correct categorization for call types of each MarV stimulus in PP (**c**) and AP (**g**) of marmoset M160E. **d**, **h** Comparisons of correct categorization between original (grouped by call types) and shuffled data of PP (**d**) and AP (**h**) for marmoset M160E (*: $p < 0.05$, rank sum test). Chance performance would be 1/4 for call types categorization (indicated by the horizontal red line in each graph).

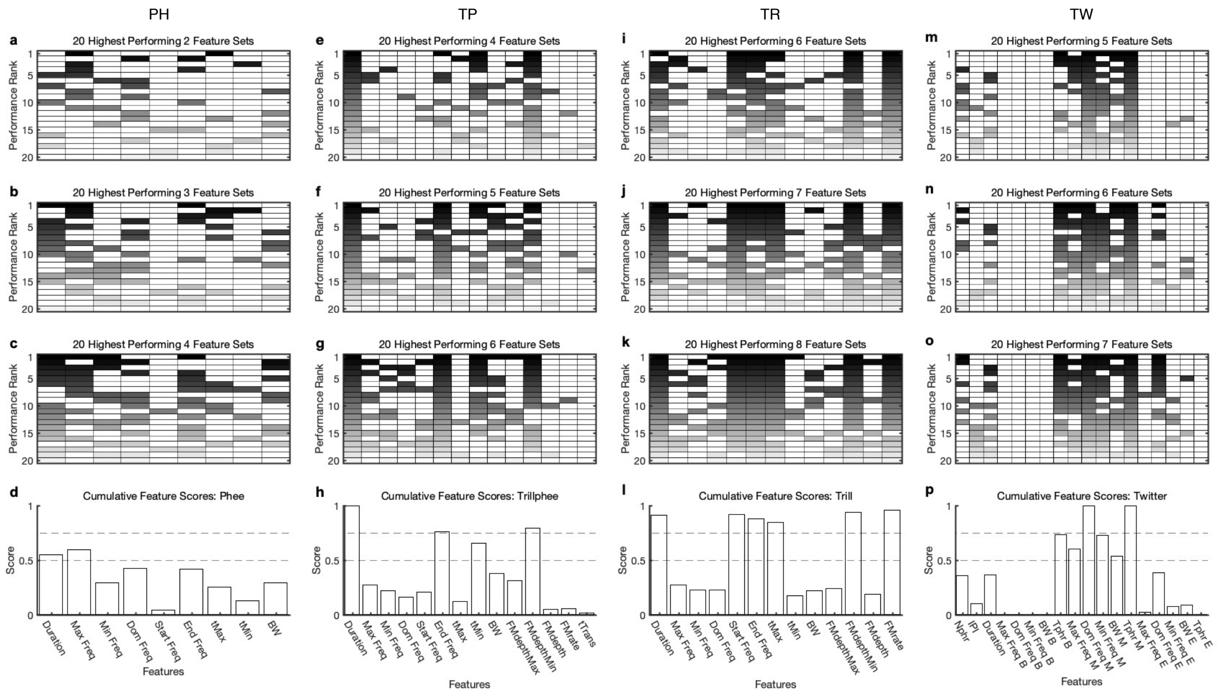

**Extended Data Fig. 8 Determine important acoustic features for identity categorization.**

**a, b, c** The best features in the top 20 performing feature sets for sets consisting of 2, 3, and 4 features for PH. The darkened boxes in each row indicate which features are included in the feature set yielding the rank-ordered performance indicated. Darker shading indicates higher-ranking feature sets and reflects the higher scores applied to each feature in the set. **d** Cumulative scores for all features used for PH. The horizontal dashed lines mark the threshold of 0.5 and 0.75. They are used to determine the notable features and critical features. **e, f, g** The best features in the top 20 performing feature sets for sets consisting of 4, 5, and 6 features for TP. **h** Cumulative scores for all features used for TP. **i, j, k** The best features in the top 20 performing feature sets for sets consisting of 6, 7, and 8 features for TR. **l** Cumulative scores for all features used for TR. **m, n, o** The best features in the top 20 performing feature sets for sets consisting of 5, 6, and 7 features for TW. **p** Cumulative scores for all features used for TW.

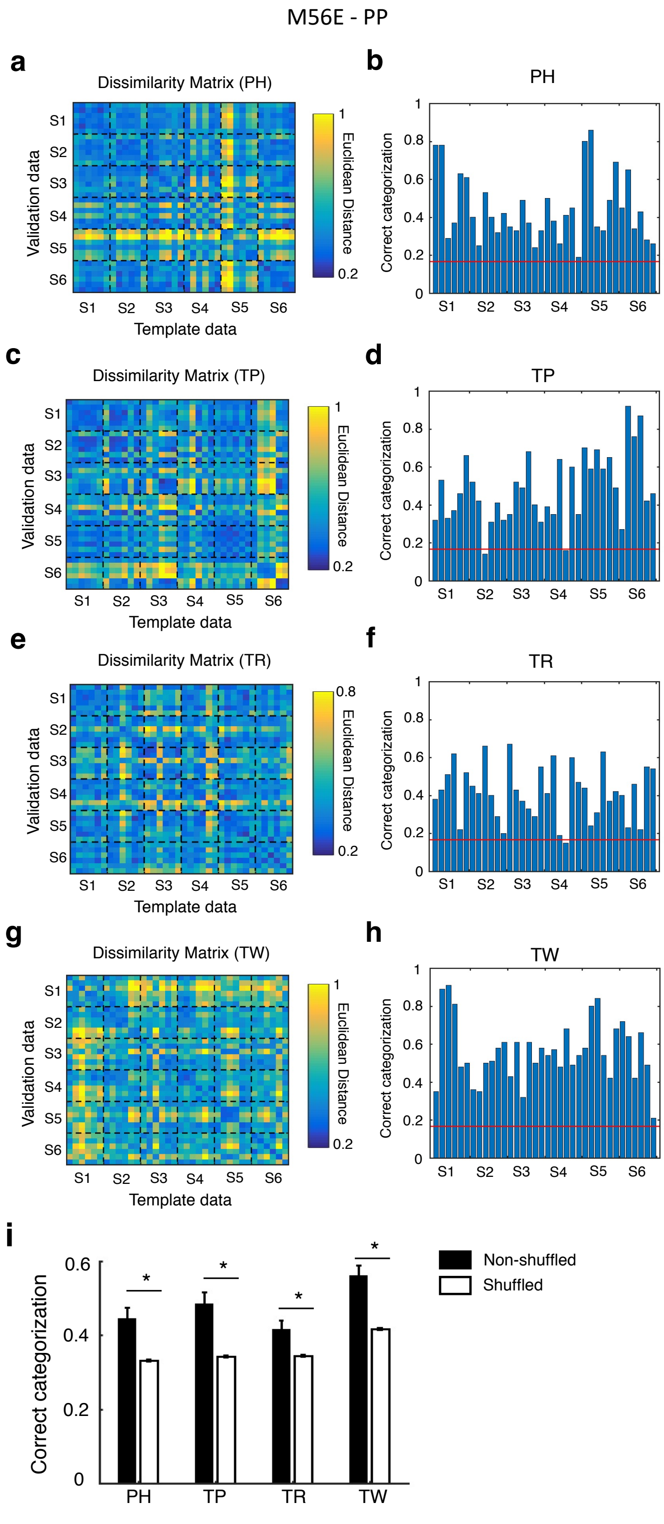


**Extended Data Fig. 9 Similar identity categorization results based on neuronal responses in PP of marmoset M56E as in AP of marmoset M56E shown in Fig. 4**

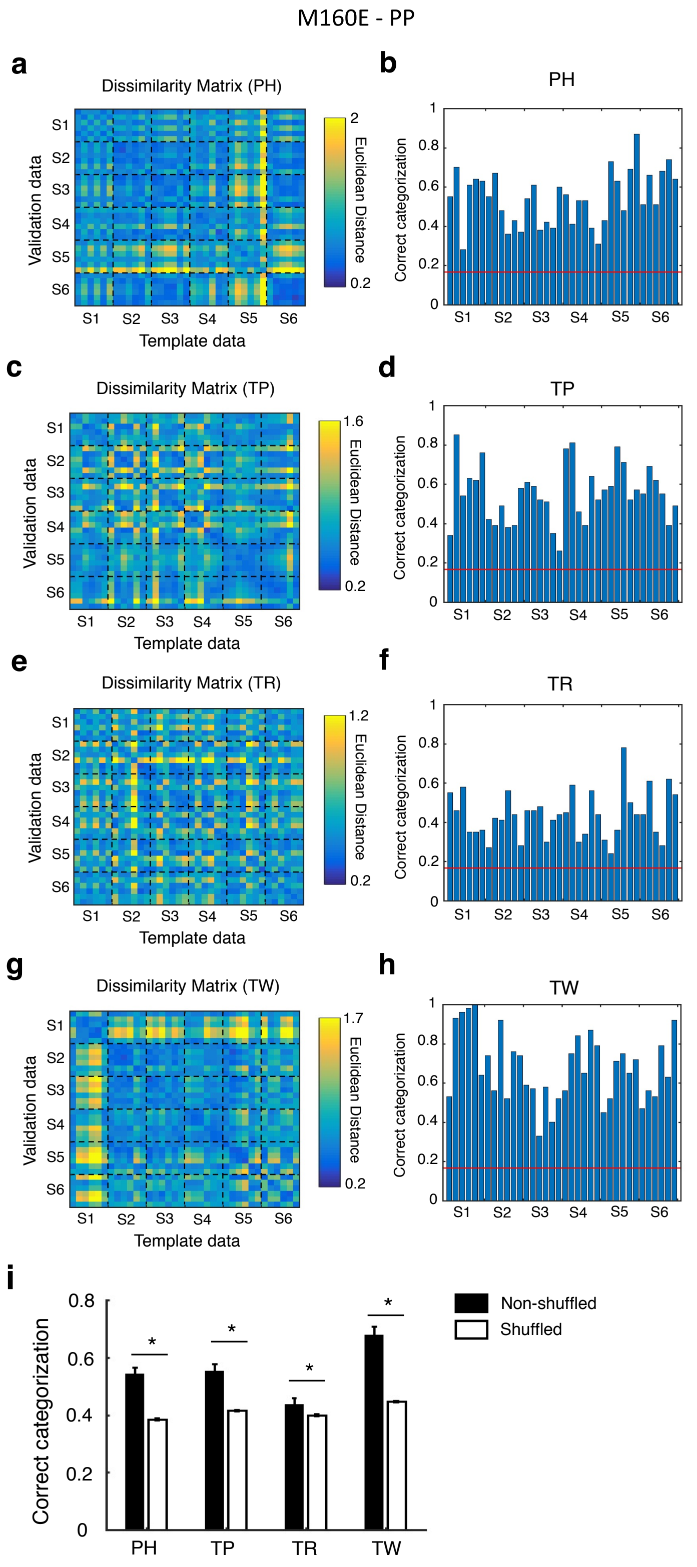


**Extended Data Fig. 10 Similar identity categorization results based on neuronal responses in PP of marmoset M160E as in AP of marmoset M56E shown in Fig. 4**

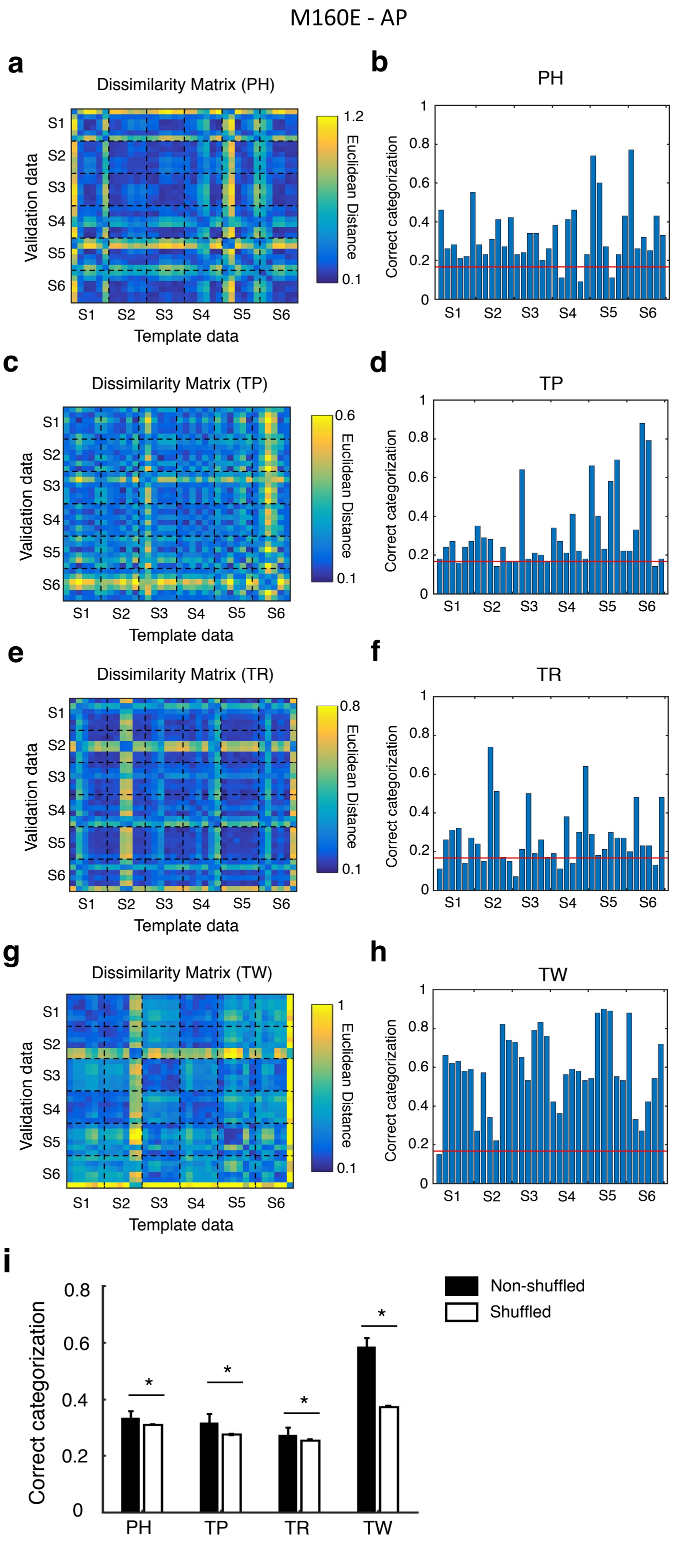


**Extended Data Fig. 11 Similar identity categorization results based on neuronal responses in AP of marmoset M160E as in AP of marmoset M56E shown in Fig. 4**
